# scCobra: Contrastive cell embedding learning with domain-adaptation for single-cell data integration and harmonization

**DOI:** 10.1101/2022.10.23.513389

**Authors:** Bowen Zhao, Dong-Qing Wei, Yi Xiong, Jun Ding

## Abstract

The rapid development of single-cell technologies has underscored the need for more effective methods in the integration and harmonization of single-cell sequencing data. The prevalent challenge of batch effects, resulting from technical and biological variations across studies, demands accurate and reliable solutions for data integration. Traditional tools often have limitations, both due to reliance on gene expression distribution assumptions and the common issue of over-correction, particularly in methods based on anchor alignments. Here we introduce scCobra, a deep neural network tool designed specifically to address these challenges. By leveraging a deep generative model that combines a contrastive neural network with domain adaptation, scCobra effectively mitigates batch effects and minimizes over-correction without depending on gene expression distribution assumptions. Additionally, scCobra enables online label transfer across datasets with batch effects, facilitating the continuous integration of new data without retraining, and offers features for batch effect simulation and advanced multi-omic batch integration. These capabilities make scCobra a versatile data integration and harmonization tool for achieving accurate and insightful biological interpretations from complex datasets.

## Introduction

Single-cell genomics technology significantly advances the exploration of complex and heterogeneous biological systems by providing high-resolution measurements of cellular states. Over recent years, the proliferation of single-cell maps detailing diverse cell types from various tissues across numerous organisms, including humans and mice, has been observed^1–5^. These comprehensive single-cell atlases are often compiled across multiple batches, necessitated by the limitations of current single-cell sequencing technologies, experimental constraints such as budget or sample availability, or the pooling of datasets from different laboratories. Additionally, the use of various single-cell sequencing platforms, like 10X Genomics^6^, micro-well^3^, and Smart-seq^7^, contributes to systematic technical variations known as “batch effects”^8–11^ between single-cell measurements across different batches. To derive a more accurate and deeper understanding of biological insights from those single-cell maps, it is imperative to mitigate batch effects among single-cell datasets and ensure their integration and harmonization^12^ for a comprehensive analysis.

To mitigate the impact of such batch effects on downstream single-cell data analytics, efficient computational methods to integrate single-cell data from different batches and remove the batch effects are essential. As of this date, dozens of single-cell data integration methods have been developed. The most widely used methods include Seurat^13^, Scanorama^14^, scVI^15^, and Harmony^16^. Seurat uses canonical correlation analysis to project the cells into a reduced latent space to identify correlations across datasets. The mutual nearest neighbors (MNNs) are then identified in the reduced latent space and serve as “anchors” to align the data from different batches. Another method Scanorama^14^ also searches for MNNs in dimensionality-reduced space and uses them to guide batch correction. Unlike the Seurat which searches similar cells across batch pairs to compute the correction, Scanorama searches across all batches and calculates the dataset merging priority based on the matching cell percentage in each batch. scVI^15^ specifically models the gene expression variance induced by the library size difference and batch effects, which will be added as additional modules to a variational autoencoder that learns the cell embeddings by minimizing the cell gene expression reconstruction error, thereby separating biological signals from batch effects. An efficient method Harmony first reduces the high-dimensional gene expression into the reduced space using principal component analysis (PCA)^17^. In the PCA reduced space, Harmony iteratively removes the present batch effects. In each iteration, the method clusters cells from different batches while maximizing the diversity of batches within each cluster and then calculates a correction factor for each cell to be integrated.

Although existing methods have demonstrated success in integrating single-cell datasets from different batches and even distinct sequencing platforms^16^, various problems and limitations remain, leaving room for further improvement. MNNs-based models find similar cells between batches and calculate the correction factor to mitigate the difference across batches. Therefore, this category of methods often over-corrects the batch effects since they tend to ignore the potential inherent difference associated with different batches, which could be problematic in many applications^18^. Specifically, depending on specific experimental scenarios, the cellular states of even the same cell types across different experimental batches could differ, which is quite common in many disease studies, in which the same cell type (e.g., Macrophages) could be significantly different between healthy and disease patients^19^. Therefore, ignoring those biological differences between batches may cripple the downstream single-cell data analytics. Deep learning-based methods such as scVI (auto-encoder based)^15^ learn the reduced cell embeddings in the latent space by reconstructing the input gene expression. It often assumes that the gene expression (i.e., raw count) follows a negative binomial distribution (or zero-inflated negative binomial)^20, 21^, but it is challenging to find a universal gene expression distribution for various single-cell datasets from different studies and by different sequencing platforms^22, 23^. A proper selection of the gene expression distribution will determine the reconstruction loss and dramatically influence the cell embedding learnings, which is critical for the downstream batch correction^24^. Finally, certain methods and their original implementations are challenging to apply in the original feature space^8, 16^, leading to a loss of model interpretability and making it difficult to understand the differences between various cellular states at the genetic level.

Eliminating batch effects not only facilitates the integration of multiple datasets, but also augments downstream tasks such as label transfer, batch generation, and multimodal batch correction. For label transfer tasks, some methods, such as Seurat and MNN, require mapping new data onto a reference atlas for cell type annotation. However, this process is computationally intensive due to the large number of cells in the reference atlas. Moreover, these methods necessitate model retraining whenever new data becomes available, limiting the efficient utilization of pre-trained models^13, 15, 16, 25^. Besides, the evaluation of current batch correction methods heavily relies on the simulation datasets^26, 27^. However, there remains a discrepancy between the simulated batch-affected single-cell data and its real single-cell data counterpart^28^. This can easily result in inaccurate evaluations and benchmarking of batch correction methods’ performance. In addition, integrating different omics data types offers a more comprehensive perspective of cellular states. While some methods have shown effectiveness in reducing batch effects in scRNA-seq data^12, 29^, their ability to incorporate scATAC-seq data and tackle batch effects across multi-omic datasets remains limited. Additionally, many current multi-omics integration methods are performed in latent space^30–34^, which is not conducive to using the mature single-cell RNA sequencing (scRNA-seq) analysis workflows and makes it difficult to understand multimodal data at the gene level.

To address the above limitations, here we introduce a deep neural network framework scCobra, which employs contrastive learning at both cellular and cluster levels, variational Autoencoder (VAE) and Generative Adversarial Networks (GANs) to integrate single-cell data across varying batches. scCobra distinguishes itself through its comprehensive approach to data integration and harmonization across single-cell studies. Its proficiency spans a range of critical functions, from effectively correcting batch effects in scRNA-seq datasets to handling multi-omic data variations and generating meticulously detailed single-cell datasets for rigorous benchmarking.

The capabilities of scCobra in batch correction have been validated across various tasks, including scRNA-seq, spatial omic, and single-cell multi-omic. Through both simulation and analyses of real disease datasets, scCobra has been shown to effectively minimize the risk of over-correction in batch adjustments. Beyond batch correction, scCobra enables a range of downstream data harmonization analyses, such as online cell label propagation and the creation of benchmark batch-affected single-cell datasets enriched. The introduction of scCobra equips the single-cell genomics research community with a tool designed to tackle batch effects, enhancing the integration and harmonization of relevant single-cell datasets and aiding in the generation of precise and reliable biological insights.

## Result

### scCobra model overview

scCobra is a deep neural network framework that employs the contrastive VAE-GAN architecture, designed for the integration and harmonization of single-cell data across multiple batches (Fig.1). Using an encoder, scCobra maps the single-cell data across different batches into a latent low-dimensional space. A decoder with a Domain Specific Batch Normalization (DSBN) layer then reconstructs the original inputs with batch effect from the latent representations. The reconstruction objective of scCobra has three primary facets. The first is an adversarial training component, ensuring that the reconstructed data closely mirrors the original. The second part is the reconstruction loss, which ensures that the batch information is accurately reconstructed. The third is a contrastive learning component, split between cell-level and cluster-level mechanisms. At the cell level, it ensures maximal similarity between original cells (view1) and their reconstructed counterparts (view2) in the latent space. Meanwhile, at the cluster level, it mandates consistency in cluster category outputs between original (view1) and generated cells (view2). A domain discriminator is also incorporated to remove batch information from the latent embedding *μ*. Our method can also deliver batch correction in the original input space by reconstructing all input cells from the batch-corrected latent space via the same BN layer (e.g., BN1). The versatility of scCobra’s batch-corrected outputs, effective in both latent and original spaces, makes it highly applicable for a variety of downstream tasks. These include batch correction, multi-omic data integration, online label transfer and the simulation of single-cell datasets with batch effects. Detailed description of the scCobra model is available in the Methods section.

**Fig. 1.**
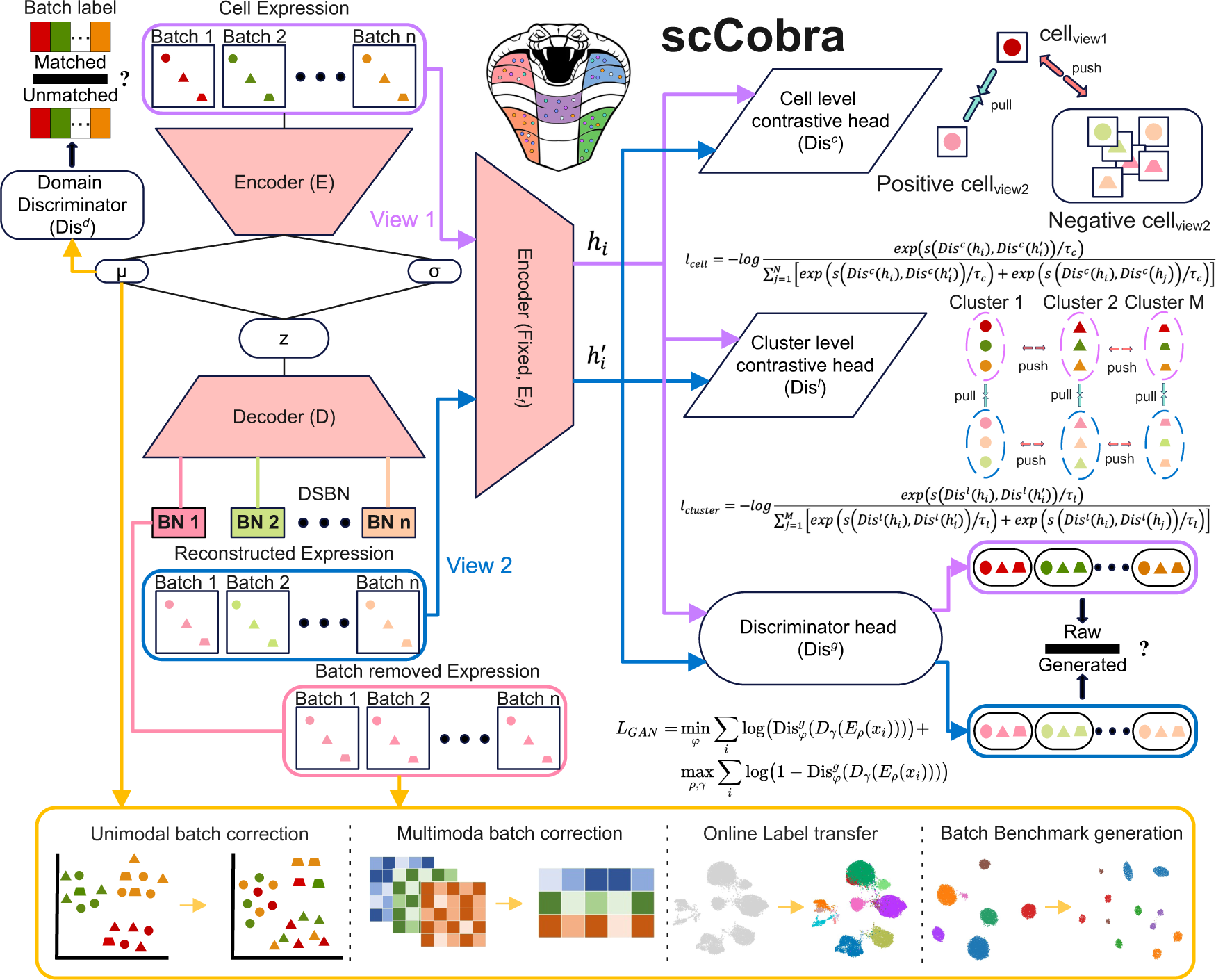
scCobra method overview. scCobra integrates a contrastive variational autoencoder with a generative adversarial network architecture to process multiple single-cell data batches. This model features an encoder that consolidates inputs from various batches into a unified low-dimensional latent space. A decoder, enhanced with a Domain-Specific Batch Normalization (DSBN) layer, then reconstructs the original batch data. The DSBN layer is specifically designed to recognize and adjust for batch effects, enabling accurate reconstruction of batch-influenced data from the latent space. The reconstruction aim of scCobra is split into three primary objectives. The first is adversarial training, focused on increasing the resemblance between the reconstructed and original data. The second part is the reconstruction loss, which ensures that the batch information is accurately reconstructed. The third leverages contrastive learning at both cell and cluster levels to ensure precision in data replication and category consistency between original and generated cells. A domain discriminator further refines the model by extracting batch-specific signals from the latent embeddings (μ), enhancing the purity of the latent space representation. The dual outputs of scCobra, from both latent (μ) and original space (Batch removed Expression, all batches mapped to BN1), support a broad spectrum of downstream applications. These include but are not limited to unimodal batch correction, multimodal batch correction, label transfer and the generation of benchmark datasets, demonstrating scCobra’s adaptability and utility in single-cell data analysis.

### scCobra mininizes over-correction risks in batch correction

The goal of batch correction is to minimize these technical variations so that the true biological variations across samples can be accurately identified and analyzed. However, over-correction occurs when the batch correction algorithm is too aggressive, leading to the loss of relevant biological information, which is a prevalent challenge among existing methods and often hinders the adoption of the batch correction in practical applications^18, 25, 35^. To showcase that scCobra can minimize the over-correction risks in comparison to its counterparts, here we applied the methods to a simulated dataset (human immune dataset ^36–38^) and a real single-cell dataset (COVID-19^39^).

First, we adjusted the gene expression levels in *CD4*+ T cells within a human immune dataset by modifying standardized expression values for genes in the “Defense response to Virus” GO term. This was performed to simulate the gene expression changes in *CD4*+ T cells following viral infection, designating this modified group as “perturbed *CD4*+ T cells” (for detailed methods, see Methods section). After applying batch correction tools like Seurat, Scanorama, Harmony, and scCobra, we produced UMAP visualizations (Fig. 2a). In the uncorrected data, perturbed *CD4*+ T cells formed a distinct cluster within the 10X batch. Post-correction, only Scanorama and scCobra were successful in distinguishing the perturbed *CD4*+ T cells from their unperturbed counterparts. Seurat and Harmony integrated both *CD4*+ T cell groups, indicating a significant overcorrection. Scanorama, while distinguishing between the two, also indicated a potential for under correction as many cells of the same type appeared scattered across different clusters. Further analysis into overcorrection scores reaffirmed these observations (Fig. 2c), with scCobra outperforming the others by maintaining clear separation between perturbed and standard *CD4*+ T cells, as evidenced by distinct expression of key marker genes like *IL27* and *TLR9* (Fig. 2b).

**Fig. 2:**
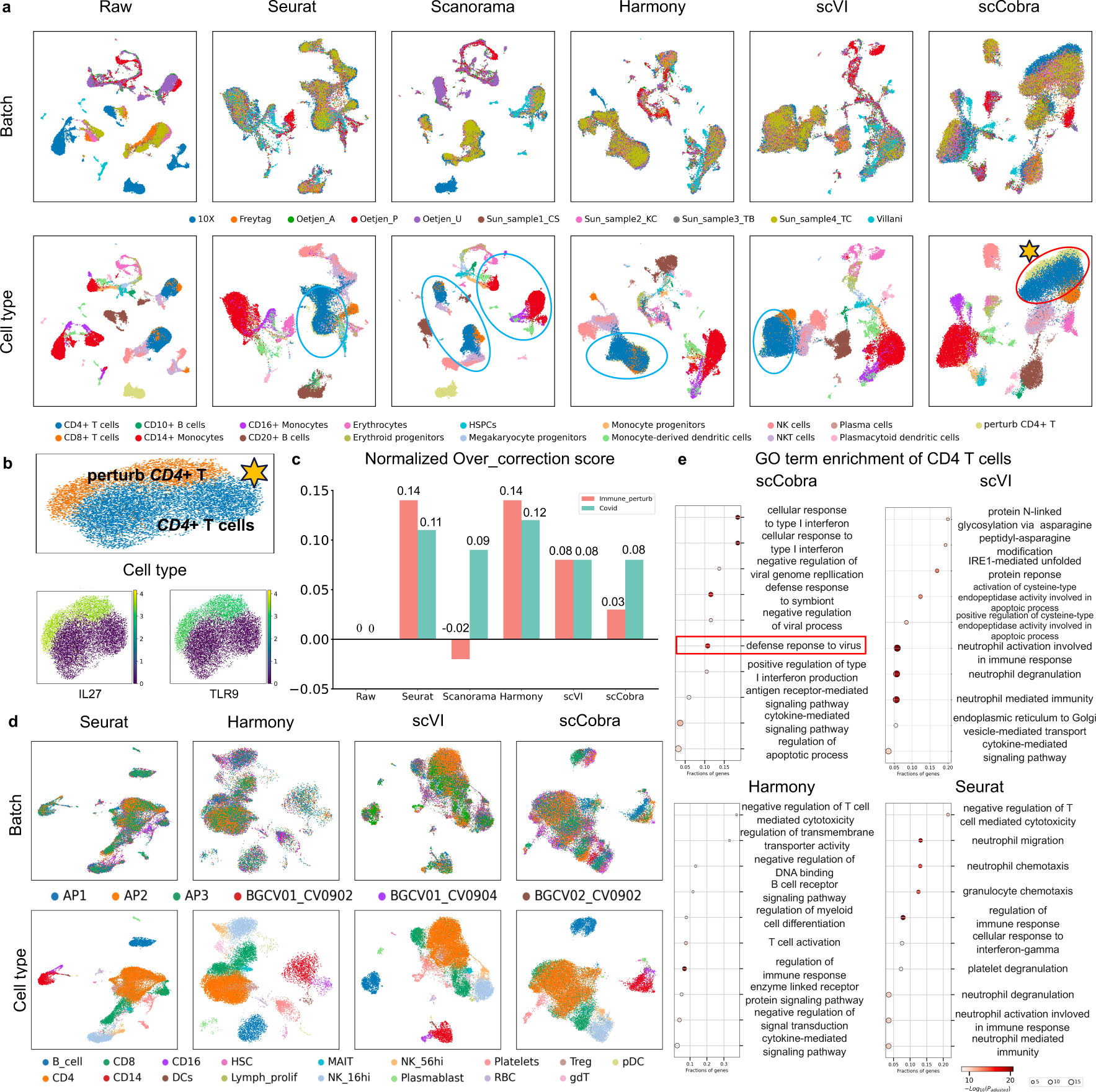
scCobra Can Minimize Over-correction Risk. **a**, Batch correction methods (Seurat, Scanorama, Harmony, scVI, scCobra) were assessed on a simulated immune single-cell dataset, introducing batch effects and biological differences by perturbing genes associated with the “Defense response to virus” Gene Ontology (GO) term. The figure’s top panels are color-coded by batch, and the bottom panels by cell type, showing each method’s effectiveness in preserving biological distinctions while correcting batch variations. Seurat, Harmony, and scVI tend to merge *CD4*+ T cells from various conditions, suggesting over-correction, whereas Scanorama often results in under-correction. In contrast, scCobra effectively differentiates between original and biologically altered *CD4*+ T cells. **b**, The enlarged UMAP plots clearly indicate that scCobra separates normal from perturbed *CD4*+ T cells, highlighted by specific markers *IL27* and *TLR9*. **c**, Quantitative analysis on both the simulated immune dataset and an independent COVID single-cell dataset indicates Seurat, Harmony, and scVI have high over-correction scores, indicative of over-correction, while Scanorama shows a negative over-correction score, signaling under-correction. scCobra maintains a balanced and accurate correction, preserving the biological variance found in the raw data. **d**, UMAP visualizations of an independent COVID-19 dataset corrected by different methods showcase their batch correction capabilities, with the first row colored by sample origin and the second by cell type, illustrating their real-world effectiveness. **e**, Post-correction Gene Ontology enrichment analysis for the COVID-19 dataset underscores scCobra’s capability to retain the “Defense response to virus” GO term, which is a top-identified term in the raw (uncorrected) COVID data, crucial for COVID sample analysis. This stands in contrast to other methods, which exhibit correction inaccuracies, evidenced by the loss of this critical GO term, indicative of over-correction and information loss.

Second, to further validate scCobra’s ability to mitigate overcorrection, we focused on real- world scenarios, using a dataset that juxtaposed three healthy samples against three from Covid-19 patients, considering each patient sample as its distinct batch. After applying Seurat, Harmony, and scCobra for batch correction, UMAP visualizations were employed (Fig. 2d). All three methodologies adeptly countered batch effects. However, an enrichment analysis conducted on all genes with *P*-values less than 0.05 revealed critical insights into the distinct expression profiles between healthy and Covid-19 *CD4*+ T cells. A subsequent GO term enrichment analysis (Fig. 2e) showed that genes from both the raw dataset (Supplementary Fig. 2) and those processed via scCobra were enriched in the “Defense response to Virus” GO term (Adjusted *P* 1.39 × 10^!”^). Significantly, scCobra’s results were more pronounced, indicating its proficiency in balancing batch correction without sacrificing biological differences. In contrast, while Seurat and Harmony did manage to counteract batch effects, their results masked the essential biological variance between the two groups, an indicator of overcorrection, this conclusion is also supported by the overcorrection score^25^ calculated through benchmark testing (Fig. 2c).

### scCobra enables multi-omic batch correction

While many contemporary methods target the correction of batch effects in single-cell RNA-seq data exclusively, scCobra stands out due to its versatility, adeptly addressing batch effects across a spectrum of single-cell data types, not just specifically applied to scRNA-seq. Here, we harnessed scCobra for single-cell multi-omic data harmonization. We transformed the scATAC-seq peak matrix into a gene activity matrix using MAESTRO^40^ and treated it like a specialized scRNA-seq batch, streamlining the integration of single-cell multi-omic data into a batch correction task. However, even after converting the scATAC-seq peak matrix into a gene activity matrix, the modal differences with scRNA-seq data remained substantial. scCobra helped mitigate this issue, the batch-corrected fusion of the two omics data showcased a remarkable alignment: not only were cells of congruent types grouped, but those sharing analogous developmental arcs also neighbored each other. For instance, B cell progenitors aligned with pre-B cells, and *CD14*+ Monocytes nestled alongside *CD16*+ Monocytes (Fig. 3a). From the perspective of quantitative metrics, Harmony and Scanorama struggle to handle multi-omic data with significant modal differences, whereas scCobra, scVI, and Seurat are all capable of effectively integrating multi-omic data (Fig. 3b). However, scCobra retains the most biological features (Fig. 3c).

**Fig. 3.**
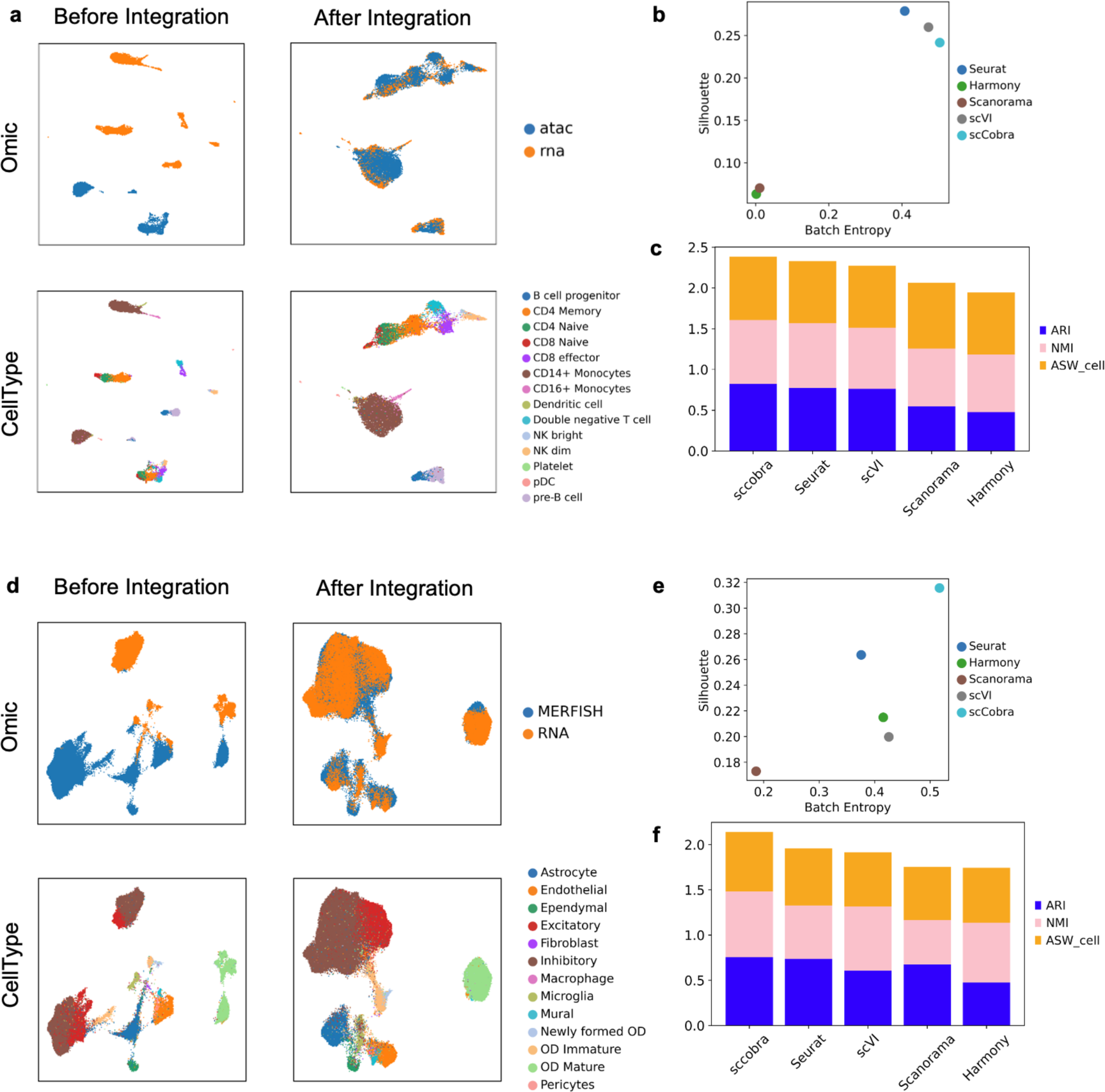
Multimodal batch correction. **a**, the UMAP visualization results before and after batch correction of single-cell multi-omic dataset. In the first row, the colors represent the different omics, while in the second row, the colors represent the cell types. **b**, **c** the dot plot and bar plot show the performance of scCobra compared to competing methods in integrating multi-omic data. **d**, the UMAP visualization results of the single-cell spatial dataset before and after batch correction. In the first row, the colors represent the MERFISH data and RNA-seq data respectively, while in the second row, the colors represent the cell types. **e**, **f** the dot plot and bar plot show the performance of scCobra compared to competing methods in integrating spatial omic data.

To further expand its applicability in data integration and harmonization, we integrated scRNA-seq data with the single-cell spatial MERFISH data^41^ (Fig. 3d). Due to the relatively fewer genes detected by MERFISH compared to scRNA-seq, it is challenging to integrate such data using a limited set of features. We selected genes expressed in both MERFISH and scRNA-seq data as inputs for testing. The results indicate that scCobra effectively integrated MERFISH data with scRNA-seq data, for instance, positioning OD Immature cells adjacent to OD mature cells. From the perspective of quantitative metrics, Scanorama struggles with handling batch effects between scRNA-seq and MERFISH data. Seurat, Harmony, and scVI have similar performances, whereas scCobra demonstrates the best performance in handling batch effects (Fig. 3e). In terms of biological conservation metrics, Harmony and Scanorama preserve a similar level of biological specificity. Seurat and scVI perform better than the first two methods, while scCobra significantly outperforms all the competing methods (Fig. 3f).

### scCobra enables simulation of single-cell data with batch effects

While many single-cell batch correction methods rely on integrating multiple datasets for performance evaluation, inherent variations in annotation standards across these datasets can induce biases, Some methods, such as Splatter^27^, can simulate batch effects by setting different random seeds, but they cannot be applied to simulated data based on a real dataset, which potentially limits the authenticity of data and its application in benchmarking batch correction methods. Addressing this challenge, scCobra can simulate batch effects with a ground-truth reference based on an existing real single-cell dataset, ensuring an unbiased evaluation of batch correction techniques. Specifically, for a given batch ‘n’, scCobra first secures an embedding devoid of batch information using a pretrained encoder. Thereafter, with its trained decoder and DSBN module, scCobra can simulate distinct batches from one original batch via different BN layers for decoding from the latent space (Fig. 4a). This approach guarantees uniform cell type annotations across the simulated batches, removing potential annotation inconsistencies.

**Fig. 4.**
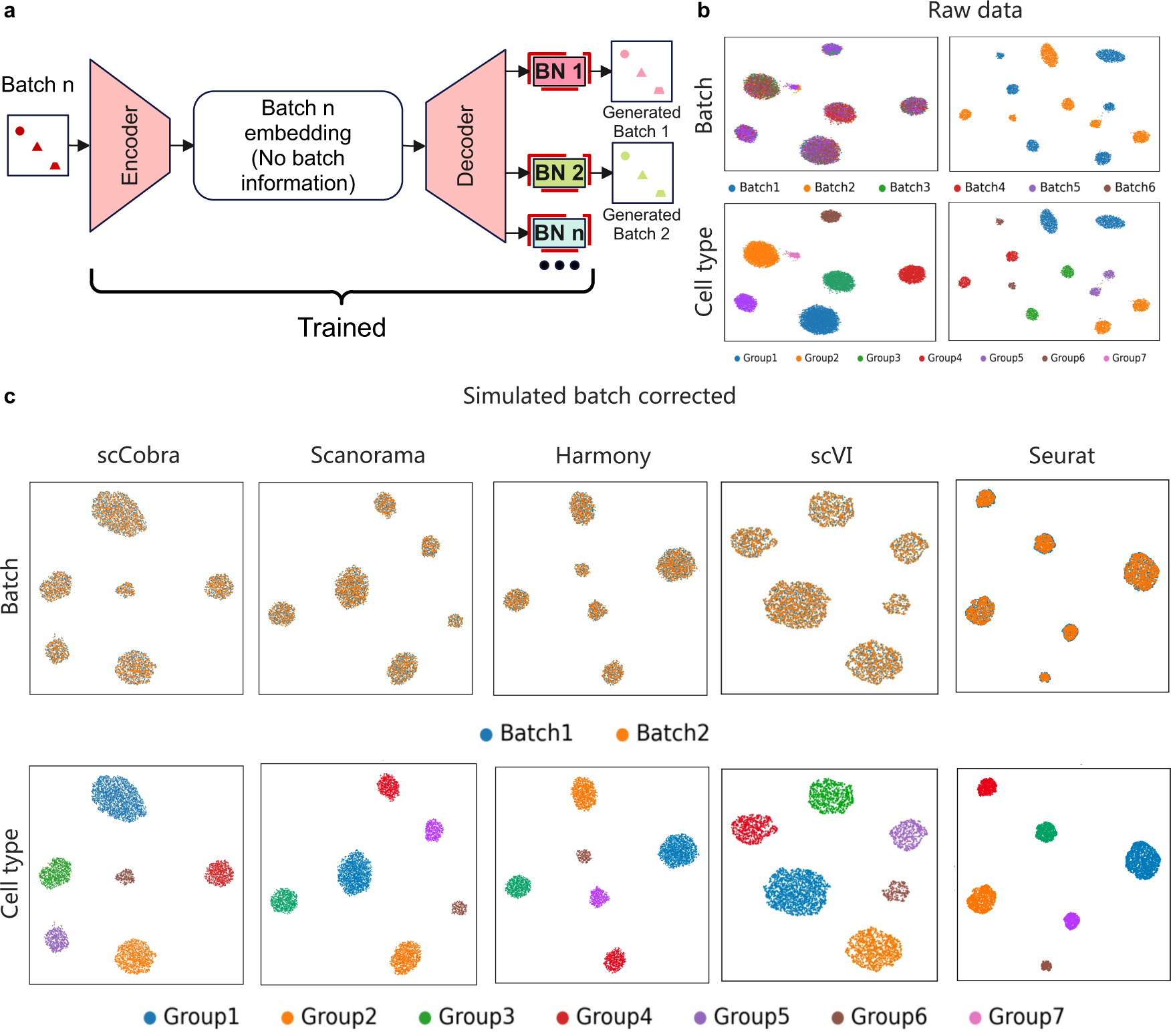
scCobra’s Efficacy in Simulating Single-Cell Data with Batch Effects. **a**, Illustration of scCobra’s capacity to generate single-cell RNA-seq data simulations that include batch effects, showcasing the adaptability and application of the scCobra model. **b**, Through UMAP visualization, this section demonstrates the effectiveness of scCobra in eliminating batch effects from data, as well as showcasing real single-cell data re-generated with batch labels, highlighting its practical utility in real-world scenarios. **c**, Features UMAP visualizations of simulated data after batch correction, with these panels designed to illustrate the efficiency of various methods in correcting batch effects within datasets simulated by scCobra. It is important to note that because the data was simulated by scCobra, its batch correction is inherently advantaged; however, this simulation acts as a benchmark to evaluate the performance of other batch correction tools such as Harmony, Scanorama, scVI, and Seurat, as shown in panel c.

To illustrate this functionality and the process of generation, we created a simulated single-cell dataset that exhibits batch effects, derived from a consistent set of cells annotated with their cell types, thereby establishing a clear ground truth. This approach was aimed at mirroring real-world batch variations while maintaining precise biological annotations for subsequent validation. Initially, we employed scCobra to address the batch effects present within this dataset, evaluating the correction’s effectiveness through visual analysis with UMAP projections (Fig. 4b). Following this, scCobra demonstrated its versatility by replicating cells from Batch 1 within both the original Batch 1 and an artificially created Batch 2, effectively generating simulated batches for comparative analysis. In the visualizations produced by Batch UMAP, we employed a color-coding scheme where blue indicated cells from Batch 1 and orange denoted those from Batch 2, facilitating an intuitive understanding of batch origins. Additionally, the cell type UMAP visualization was leveraged to distinctly categorize the cell types (Fig. 4c). Building on this foundation, we broadened our analysis to include the application of scCobra, Scanorama, Harmony, scVI and Seurat to this intentionally simulated dataset, aiming to scrutinize the batch integration efficacy of each method. All three methodologies proficiently facilitated the amalgamation of cell populations from both batches, as clearly depicted in the cell type UMAP, where cells of identical types were aggregated (Fig. 4d, e, f).

### scCobra provides a flexible online label-transfer framework

Label transfer is a crucial data harmonization task and often requires consideration of batch effects between unlabeled data and reference datasets. Therefore, many conventional methods^13, 15, 16^ first perform batch correction on the reference atlas and unlabeled dataset, and then employ classification methods such as KNN to transfer the labels from the reference dataset to the unlabeled dataset. Consequently, retraining is necessary each time a new dataset needs to be annotated, resulting in repetitive consumption of computational resources. In the case of annotating multiple new datasets, the computational burden is further increased. To address this limitation, scCobra can directly infer unlabelled datasets without the need to retrain the model to integrate them with the reference atlas. The saved model weights are then used to annotate new source datasets. We performed batch on the Pancreas scRNA-seq dataset, and subsequently utilized the corrected results as a reference atlas to reannotate the GSE114297, GSE8547, and GSE8339 datasets. All three datasets were mapped onto the reference atlas constructed by scCobra (Fig. 5a). In addition to the comparison with SCALEX^25^, we also utilized VAE-based method scVI and transformer- based method TOSCIA to annotate the query dataset. scVI effectively disentangles batch information and true expression values in the raw expression data. We performed joint training of the raw data from the pancreas reference dataset and query dataset, facilitating the transfer of labels from the pancreas reference dataset to the query datasets. TOSICA^42^ employs gene pathways as a mediator to directly learn the relationship between gene expression levels and corresponding cell types. We employed a pre-trained TOSICA model, trained on the pancreas reference dataset, to directly infer cell type labels for the query dataset. Notably, both scCobra and SCALEX achieved a weighted F1 score of 0.95, while TOSICA achieved 0.94. In contrast, scVI attained a weighted F1 score of only 0.92. These findings underscore the exceptional online annotation capability of scCobra.

**Fig. 5.**
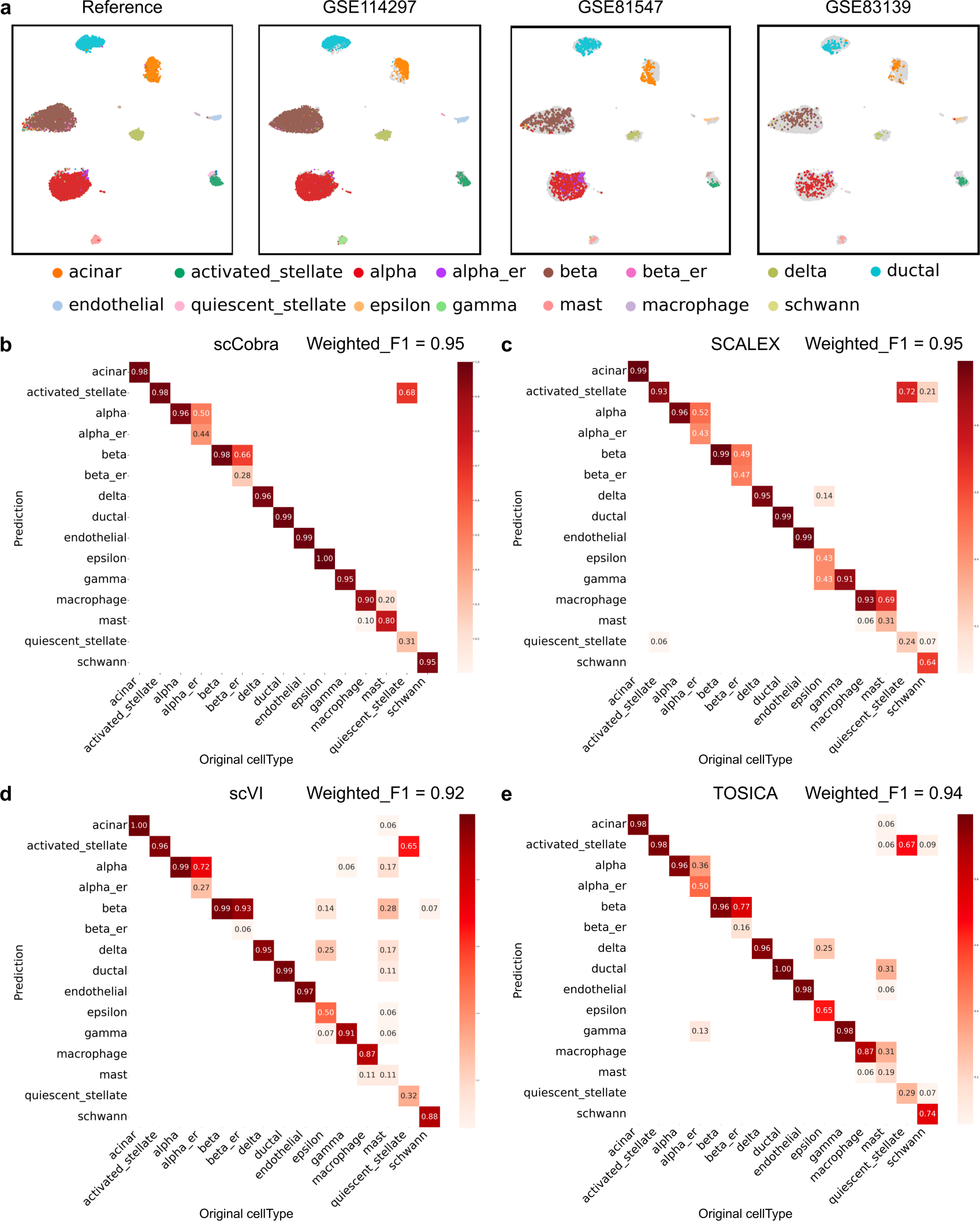
Label Transfer for Data Harmonization Across Three Different Studies Using scCobra. **a**, Features UMAP visualization results: the left side shows the reference atlas constructed using scCobra, while the right side displays integrated data from three different studies (GSE114297, GSE85447, and GSE83139), mapped onto the reference atlas and colored by cell type. This illustrates scCobra’s capability in aligning datasets from distinct studies through batch effect removal for consistent label transfer. **b-e**, present the label transfer results for scCobra (b), SCALEX (c), scVI (d), and TOSICA (e), comparing the effectiveness of each method in transferring labels from the reference atlas to the unlabeled datasets derived from the three studies. The color intensity in each panel reflects the annotation accuracy, with darker colors indicating higher accuracy. This comparison highlights the utility of scCobra in facilitating online label transfer and data harmonization across various single-cell datasets from different studies.

### scCobra outperforms benchmarking methods in batch effect correction

We employed the scCobra method on several datasets: A simulated dataset generated by Splatter^27^, a pancreas single-cell dataset encompassing data from different sequencing platforms^43–47^, and an immune cell bone marrow dataset^48–52^. To benchmark the performance and evaluate the effectiveness of scCobra, we also utilized several state-of-the-art single-cell batch correction methods, namely Seurat, Scanorama, scVI, and Harmony, on these datasets for comparative analysis.

For the simulated single-cell benchmarking dataset, the UMAP plots (Fig. 6a) highlighted a notable difference in how various methods addressed batch effects. MNNs-based approaches notably over-corrected, leading to detrimental impacts on subsequent cell clustering and cell type identification processes. For instance, while raw data distinctly separated Group 2 from Group 7 without any overlap, Seurat, Scanorama, and Harmony inadvertently merged Group 7 with Group 2. Such erroneous integrations can adversely affect downstream clustering and differential expression marker selection. Conversely, both scVI and scCobra exhibited resilience against over-correction, successfully maintaining the distinction between Group 2 and Group 7. Additionally, Seurat struggled to seamlessly integrate batch 4 and batch 6. However, the most striking evidence of scCobra’s efficacy was evident in the biological feature preservation and batch removal metrics (Fig. 6c). Here, scCobra consistently surpassed other benchmarking methods, solidifying its position as a potent tool for managing batch effects.

**Fig. 6.**
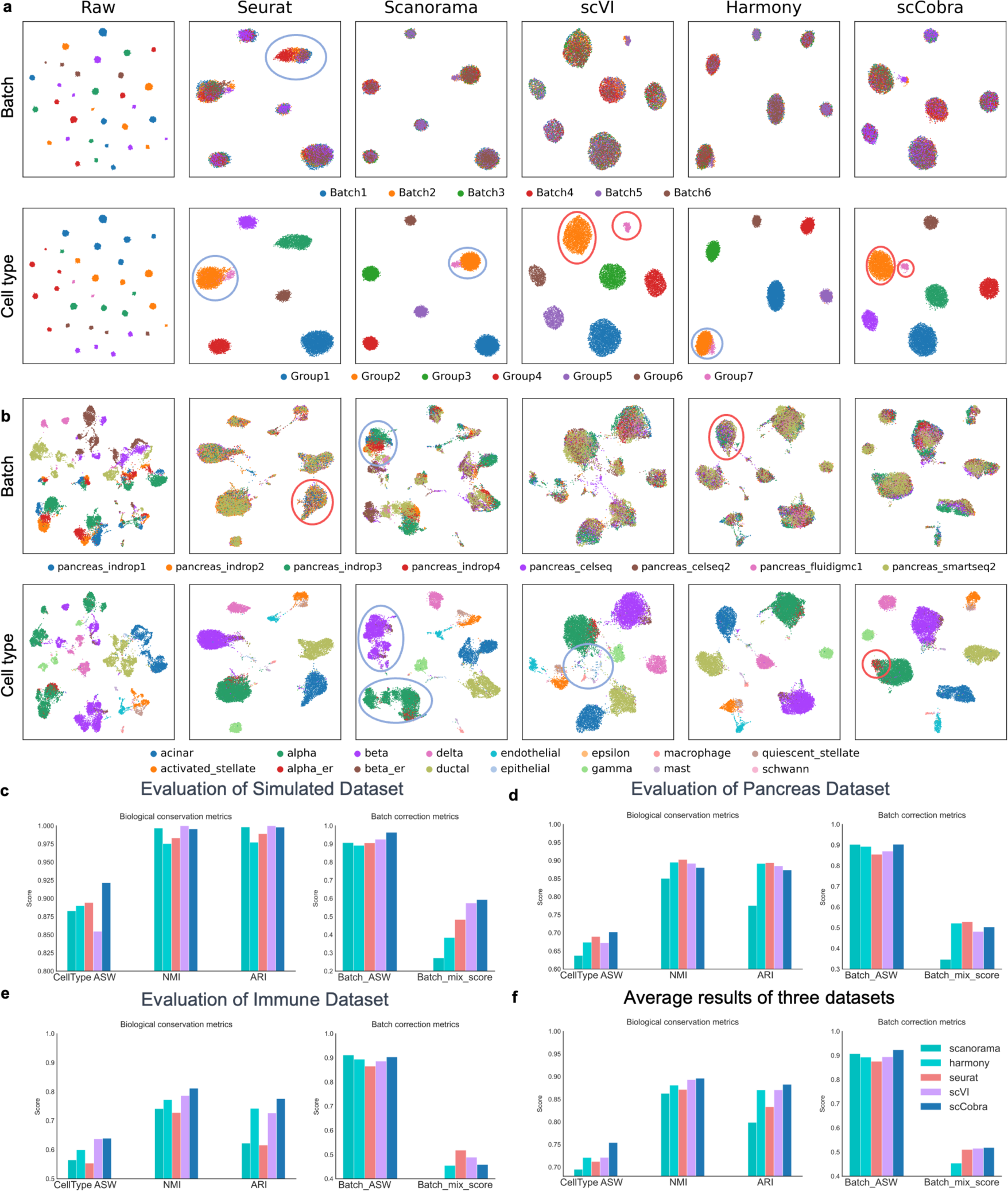
scCobra demonstrates superior batch correction performance over state-of-the-art methods. **a,** UMAP visualization comparing scCobra’s performance against benchmarking methods (Seurat, Scanorama, scVI, and Harmony) on the simulated dataset. The visualization is split into two rows: the top row shows batch correction results, with different batches distinguished by color, and the bottom row presents cell type aggregation, with each cell type assigned a unique color. Superior performance areas are circled in red, indicating where scCobra or the benchmarking method excels, while blue circles denote areas needing improvement. **b,** UMAP visualization for the pancreas dataset, comparing the batch correction capabilities of scCobra to those of benchmarking methods. Similar to panel a, the first row highlights the effectiveness of batch correction, colored by batch, and the second row shows the aggregation of cell types, with distinct colors for each cell type. **c-e,** Quantitative results of scCobra and the benchmarking methods across different datasets. This panel differentiates the performance of each method using color coding, providing a clear comparison of their effectiveness in addressing batch effects and enhancing data comparability. **f,** Average quantitative results of scCobra and benchmarking methods across different datasets, displaying the robust batch correction performance of scCobra.

Batch effects stemming from differences in single-cell sequencing platforms present a notable challenge in integrating data. Given this, we further assessed scCobra’s capability to harmonize single-cell data from varying sequencing platforms using a pancreas single- cell dataset^53^. This dataset encompasses cells derived from six distinct sequencing platforms (CEL-seq^54^, CEL-seq2^55^, Fluidigm c1^56^, InDrop^57^, SMARTer^58^, and Smart- seq2^59^) spread across nine batches. The UMAP visualization (Fig. 6b) provides insightful comparative results. Scanorama exhibited a limited correction capacity, evident from the discernible batch effects and the suboptimal integration of Alpha and Beta cells. Meanwhile, scVI’s output showed an intermingling of Macrophage cells, mast cells, and a subset of acinar cells, compromising the clarity needed to distinguish these cell types without specific markers. Contrastingly, scCobra emerged as the only method adeptly clustering alpha_er cells without confusion, whereas other tested methods showed a propensity to conflate alpha cells with alpha_er cells. In addition, scCobra’s UMAP plots were notably distinct, enabling clear delineation between various cell types, a characteristic immensely valuable for accurate clustering. Furthermore, benchmark metrics underscore scCobra’s solid performance over other benchmarking methods. It tops the charts in CellTypeASW, BatchASW, and Batch_mix_score. Moreover, scCobra’s NMI and ARI metrics are competitive, closely mirroring the results of recognized methods like Seurat and Harmony (Fig. 6d). Collectively, these outcomes attest to scCobra’s prowess in eradicating batch effects from single-cell data sourced from diverse platforms The inherent complexity of the immune dataset^53^, featuring 32,472 cells across 15,148 genes and spread over 16 batches and 17 distinctive cell types, provides an ideal canvas to assess the robustness of various integration methods in challenging scenarios. Upon examining the UMAP visualization, it’s evident that Seurat faces difficulties, particularly in effectively amalgamating batch Oetjen_P and batch Oetjen_U (see Supplementary Fig. 1). Similarly, Scanorama, despite its merits, showcases its limitations in such intricate environments. It doesn’t only grapple with the unification of cells from varied datasets but also stumbles in seamlessly integrating subsets like the *CD4*+ T cells and *CD14*+ Monocytes. scVI and Harmony, although performing commendably under less convoluted conditions, reveal their vulnerabilities within this complex framework. scVI, while integrating batches commendably, somewhat blurs the cell type boundaries, a concern that might compromise subsequent analyses. Harmony’s approach, on the other hand, leans towards over-correction, evidenced by its erroneous clustering of *CD20*+ B cells with Plasmacytoid dendritic cells. In this challenging arena, scCobra distinctly stands out. The benchmark metrics not only underscore its solid batch correction capabilities but also laud its precision in clustering that echoes the ground truth labels, particularly in such intricate datasets (as illustrated in Fig.6e). Delving deeper into the benchmark performance (Fig. 6f), while scVI registers a middle-of-the-road performance, scCobra truly excels. It emerges as the unparalleled choice for both preserving biological nuances and negating batch effects, especially when navigating the treacherous waters of complex scRNA-seq datasets.

### The contrastive learning head, discriminator head and adaBN are all indispensable for the remarkable batch correction performance

We conducted ablation experiments to dissect the key components of scCobra, examining how the removel of specific modules—contrastive learning, adversarial learning, and adaptive Batch Normalization (adaBN)—impacts its overall performance. These modified versions of scCobra were tested for batch effect correction on both Pancreas scRNA-seq and Brain scATAC-seq datasets. UMAP visualizations revealed that removing either the contrastive or adversarial learning modules significantly reduced the efficacy in correcting batch effects within the Pancreas scRNA-seq dataset. Conversely, omitting the adaBN module yielded batch correction visualizations on par with the complete scCobra model (Fig. 7a). On the Brain scATAC-seq dataset, models without contrastive or adversarial learning were unable to effectively correct batch effects. Interestingly, the model without adaBN not only differentiated between Cerebellar Granule Cells and Excitatory Neurons but also surpassed the full scCobra model’s performance in doing so (Fig. 7b). Further, we assessed the batch correction effectiveness of these four configurations using quantitative metrics. On the Pancreas scRNA-seq dataset, the scCobra model consistently excelled across various metrics (Fig. 7c). Yet, in analyzing the Brain scATAC-seq dataset, while the complete scCobra setup outperformed others in BatchASW and Batch_mix_score, highlighting its superior batch correction capacity, it slightly lagged behind the adaBN- excluded model in terms of NMI and ARI scores (Fig. 7d). This suggests that, in terms of preserving biological characteristics, the contribution of adaBN is not substantial.

**Fig. 7.**
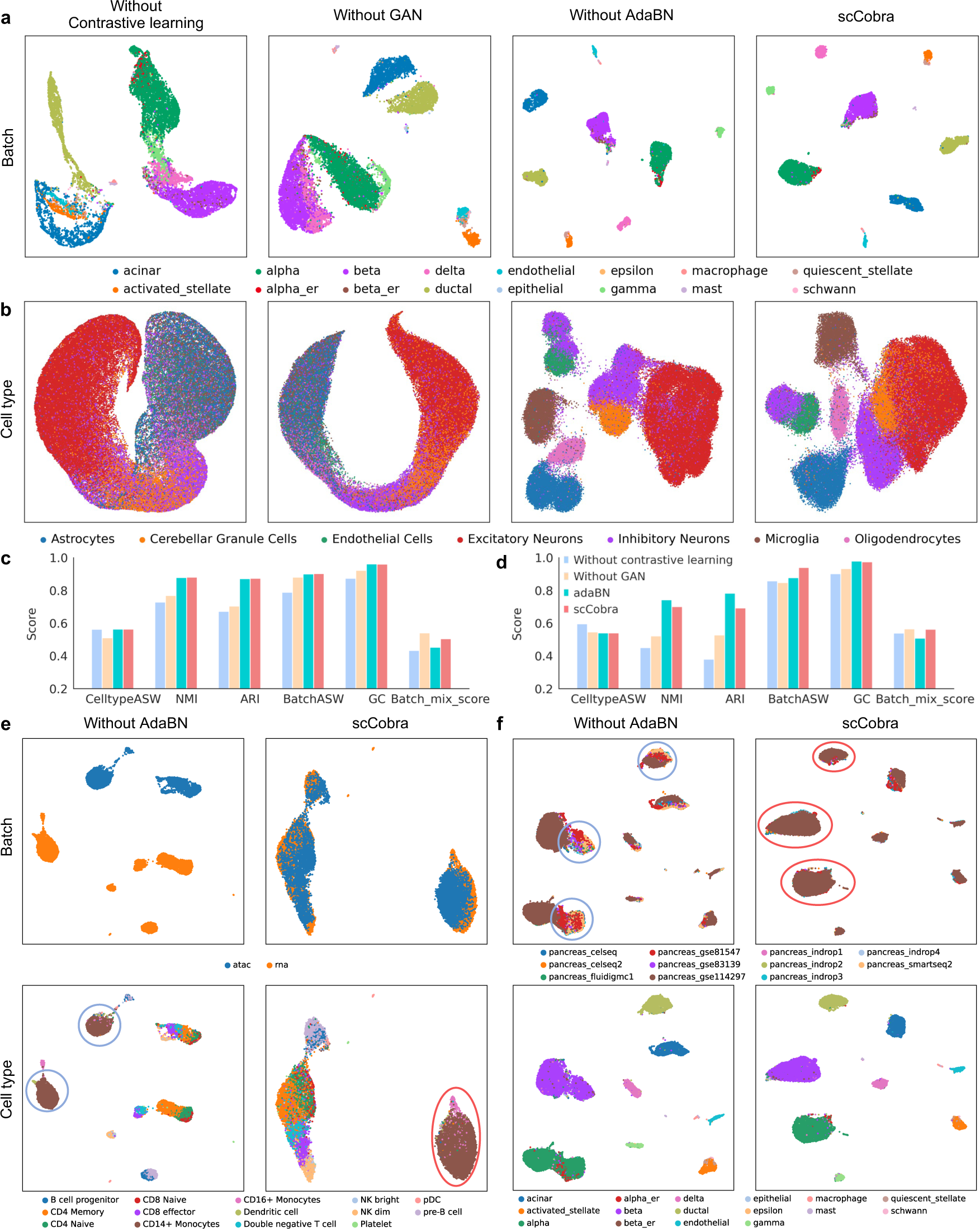
Ablation results for all components. **a**, **b** UMAP visualization results of model correction pancreas scRNA-seq dataset (**a**) and brain scATAC-seq dataset (**b**) excluding contrastive learning, excluding GAN, excluding adaBN and complete scCobra, colored by cellType. **c**, **d** quantification results of 3 ablation models and scCobra integrated pancreas scRNA-seq data set (**c**) and brain scATAC-seq dataset (**d**), colored by model. **e,** The results of integrating multi-omic data by the model without adaBN, colored by Batch and CellType respectively. Models without AdaBN failed to integrate *CD14+* Monocytes, which are highlighted with a blue circle. In the scCobra model, *CD14+* Monocytes have been integrated and are marked with a red circle. **f**, The result of label transfer for a model that does not include adaBN, colored by Batch and CellType respectively. In the UMAP plot depicting batch effects, models without AdaBN were unable to uniformly integrate data from multiple batches, as indicated by the blue circle, whereas the scCobra model achieved a more uniform integration of batches, which is highlighted with a red circle.

To examine the impact of adaBN from different angles, we carried out further ablation experiments, this time focusing on aligning single-cell datasets from varied origins. These experiments revealed that without adaBN, the model struggled to merge multi-omic data, as indicated by the failure to blend RNA and ATAC signals. On the other hand, the complete scCobra model, incorporating adaBN, successfully addressed batch effects in multi-omic datasets (Fig. 7e). Additionally, adaBN emerged as critical for effective label transfer in scCobra, enhancing its utility. While the model without adaBN could cluster cells of the same type, it was less effective in accurately relocating cells from the GSE81547 and GSE114297 batches back to their original reference datasets (Fig. 7f). These findings highlight adaBN’s vital role in dynamically aligning data from multiple sources into a cohesive space, an ability that proves essential as differences across datasets become more pronounced.

## Discussion

In this study, we developed a deep neural network based single-cell batch correction and data integration method that employs contrastive learning and generative adversarial network, which outperforms other state-of-the-art methods in single-cell data batch correction in terms of multiple batch correction evaluation metrics. We applied scCobra in 5 different single-cell datasets that represent distinct single-cell batch correction tasks, and the scCobra batch-corrected single-cell data preserve the biological information while the batch effects are minimized (as documented by the biological conservation and batch correction metrics that we benchmarked).

scCobra utilizes a deep neural network framework that integrates contrastive learning into the VAE-GAN architecture. This is designed to address the common challenges in single-cell data integration and harmonization, particularly those arising from batch effects. Here we outline the key contributions of scCobra below. (1) Over-correction risk often impedes the effective adaptation of batch correction methods and can weaken the biological signal after correction, making it challenging to discern true biological variations from artifacts. scCobra exhibits a significantly reduced risk of overcorrection compared to MNN-based methods, which is crucial for preserving the integrity of biological signals. This lower risk enhances scCobra’s ability to accurately distinguish between different disease states and identify authentic differential genes, ensuring that vital biological insights are maintained and not obscured during the batch correction process. (2) scCobra effectively addresses complex batch effects in multimodal data, a challenge where many batch correction methods faced. Unlike addressing batch effects in multi-batch scRNA-seq, the disparities within multi-omic data are considerably more pronounced. We applied a modality conversion strategy to transform scATAC-seq data into a gene activity matrix, and reframed the issue of multimodal data integration as a multi-batch integration problem.

This approach has potential benefits, by converting data from other modalities into scRNA-seq, it means that we can utilize existing mature single-cell analysis pipelines^13, 60^ to handle the data in a unified manner and conduct downstream analyses^61–63^. On the other hand, methods such as scGlue^30^ that directly integrate multimodal data in the latent space are relatively more challenging when it comes to understanding multimodal data at the gene level. Demonstrations of scCobra’s capability show it adeptly managing batch effects in tasks that integrate scRNA-seq with scATAC-seq, and in those combining scRNA-seq with spatial data, underscoring its robustness and versatility in handling diverse data types. (3) scCobra’s online label transfer functionality enables the integration and annotation of new data batches without the necessity for model retraining. This feature contrasts with many models that require joint training with a reference dataset, a process that can consume substantial computational resources. By bypassing the need for retraining, scCobra streamlines the data harmonization process essential for various downstream tasks, thus facilitating the efficient analysis of large-scale single-cell datasets. (4) Unlike other deep neural network-based methods such as scVI^15^ and scANVI^64^ which operate under specific prior assumptions regarding the gene expression distribution (e.g., negative binomial or zero-inflated negative binomial), scCobra, employing contrastive learning within its framework, does not necessitate any preconceived notions about gene expression distribution. This flexibility is particularly advantageous, addressing the challenge of finding a universally applicable gene expression distribution for diverse application scenarios. (5) Unlike the Splatter^65^, which simulates batch effects through multiple sampling, scCobra uses a DSBN layer to learn batch information from real data and adds the learned batch information to scRNA-seq data from the same batch to generate data with real batch effects. This means that scCobra understands and models the batch effects in real data, making it more suitable for generating test datasets to evaluate the batch removing capabilities of benchmark methods.

While scCobra presents notable advantages, it is not without limitations. One such limitation is that scCobra’s approach to multimodal integration relies on converting scATAC-seq peak matrices into gene activity matrices, which is contingent upon the algorithm used for this conversion process and may potentially lead to information loss, some strategies capable of performing omics transformation may be helpful, such as scCross^36^, BABEL^66^ and scButterfly^67^. Addressing this limitation could be a focus for future work, aiming to enhance the tool’s performance and versatility.

scCobra is tailored to tackle multiple challenges in integrating and harmonizing single-cell data, especially those arising from batch effects. Its ability to seamlessly and precisely integrate heterogeneous single-cell data enhances the comprehension of intricate biological mechanisms and varied cellular conditions, thereby reducing obstacles to integrating, analyzing, and leveraging large-scale single-cell datasets.

## Methods

### Data preprocessing

In preprocessing our scRNA-seq data, we first filtered out cells that expressed fewer than 200 genes and genes that were detected in fewer than 3 cells. Additionally, we removed cells where mitochondrial genes accounted for more than 5% of the total gene expression. The normalization of this filtered dataset was conducted using Scanpy’s normalize function, which standardized library sizes across the samples. This was followed by a log1p transformation to stabilize the variance. To further refine our analysis and focus on the most informative features, we identified 2000 highly variable genes for downstream analysis.

### The scCobra pipeline

The scCobra pipeline incorporates the VAE-GAN architecture along with contrastive learning and DSBN unfolding in three phases that are optimized during the training.

**Phase 1:** The domain discriminator (*Dis^d^*) is a batch label discriminator that determines whether the output of the encoder contains batch information. Its optimization goal is defined as:

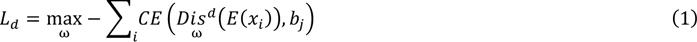

Where *E* represents the encoder, *Dis^d^* represents the batch label discriminator, *x*_i_ represents a single cell, *b*_j_ represents the batch label, *CE* represents the cross-entropy loss.

The generative discriminator is similar to the conventional GAN discriminator, which can differentiate between generated and real data. Additionally, the batch mapping network is used to simulate the original batch information within the DSBN module of the decoder. To ensure consistency between the re-introduced batch information and the original batch information, the generative discriminator is required to retain the original batch information. Therefore, the optimization goal defined as follows:

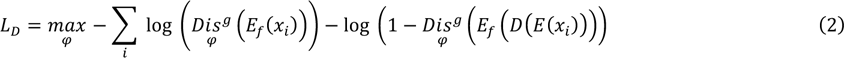

Where *Dis^g^* represents the generative discriminator, *E* represents the encoder, *D* represents the decoder, *E_f_*_’_ represent the fixed encoder (its parameters are consistent with the encoder), *x*_i_ represents a single cell, *b*_j_ represents the batch label.

**Phase 2:** After the optimization of the discriminator, we can train the generator in a GAN-like manner. Specifically, we maximize the previously trained discriminators. Since data reconstruction in the VAE-GAN framework is generated through the encoder and decoder, this step also optimizes the encoder and decoder. The optimization objective is defined as follows:

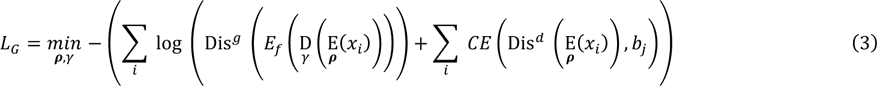

Where *Dis^g^* represents the previously trained generative discriminator from Phase 1. *E* represents the encoder, *D* represents the decoder, *Dis^d^* represents the batch label discriminator, *x*_i_represents a single cell, *b*_j_ represents the batch label, *CE* represents the cross-entropy loss.

**Phase 3:** In order to maintain consistency between the original input and reconstructed output, cell-level alignment and cluster-level alignment is necessary. To achieve this, we optimize using contrastive learning loss.

To find a good contrastive learning space, we use the VAE encoder (*E*_f’_, fixed parameters) to map both the original data and the reconstructed data into a latent space, then the output of *Cell_i_* (original cell’s expression, *x*_i_) and *Cell_i_* (reconstructed cell’s expression, *x*_i_) would be:

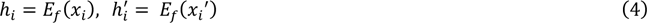

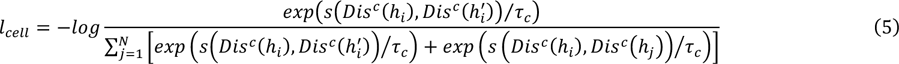

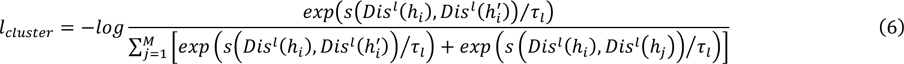

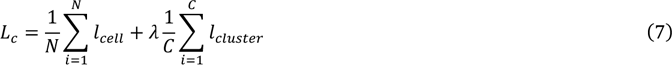

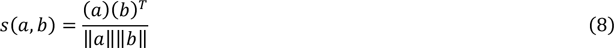

Where *l*_cell_ represents the cell level contrastive loss of the original sample, *l_cluster_* represents the cluster level contrastive loss, *L*_c_ represents the sum loss of contrastive learning, *s*(*a*, *b*) represents the cosine similarity between two vectors, *Dis*^c^ and *Dis^l^* represents the cell level contrastive head and cluster level contrastive head respectively, τ*_c_* and τ*_l_* represents the temperature coefficient of cell level contrastive head and cluster level contrastive head respectively, λ represents a weight coefficient.

In addition, we also need to add KL loss and reconstruction loss for constraints, which is defined as follows:

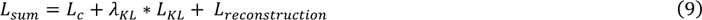

After completing the three-step optimization process described above, a full optimization cycle is achieved, and scCobra completes the optimization after multiple rounds of iteration.

### Over correction evaluation

We utilized an immune dataset to simulate a scenario aimed at examining the risk of over- correction. This involved retrieving genes associated with the “Defense of Virus” GO term from NCBI and converting them for analysis. After normalizing the dataset and applying a log1p transformation, we focused on 2000 highly variable genes. We isolated *CD4*+ T cells and increased the expression value by 1 for every gene in the “Defense of Virus” GO term within this subset. This process created a set of perturbed *CD4*+ T cells, distinct from the original cells, enabling us to assess the effects of potential over-correction in the context of enhanced viral defense gene expression. The over correction score^25^ essentially represents the complement of the frequency at which a cell and its *k*-nearest neighbors belong to the same cell type.

To compare the over-correction of scCobra and benchmark methods when integrating multi-batch datasets in the original space, we integrated three scRNA-seq datasets from COVID-19 patients with three datasets from healthy individuals, and completed the integration in the original space. To verify whether over-correction occurred, the failure to correctly enrich COVID-19-related GO term indicates that over-correction did take place. To elaborate, after completing the integration, we utilized Scanpy to select the top 200 significantly differentially expressed genes in *CD4*+ T cells between Covid-19 patients and healthy individuals. Subsequently, we performed Gene Ontology (GO) enrichment analysis using omicverse^68^, revealing the top 10 enriched GO term.

### Online label transfer

scCobra needs to be trained using single-cell datasets from multiple different batches that have been annotated to build a reference atlas. It should also save the list of input genes and the representation of each cell in the reference atlas, denoted as *emb_ref_* After the training is completed, the saved list of genes should be used to filter genes in the dataset to be annotated, ensuring that its input features match those of the training data. Next, the trained model is used to infer the cell representations in the latent space of the dataset to be annotated, denoted as *emb_querry_* In order to transfer existing labels to the dataset to be annotated, it is first necessary to train a KNN classifier using the cell representations *emb_querry_* from the reference atlas and the real labels *label_real_* Once the KNN classifier is trained, it can be used to predict the labels *label_predict_* of the cells to be annotated, thus transferring the labels from the reference atlas to the dataset to be annotated without the need to retrain the model.

We computed the label transfer accuracy individually for each cell type and normalized it based on the number of cell types, filtering out misclassified points less than 0.05. Then the heatmap was generated using seaborn. heatmap to visually represent the data. Owing to the varying quantities of cells across different types, we employ the weighted F1 score^69^ to quantify the accuracy of label annotations.

### Multimodal batch correction

In the multi-omic data preprocessing stage, we converted the open chromatin profiles from scATAC-seq into a gene activity matrix utilizing the MAESTRO package. We treated the gene activity matrix and the original scRNA-seq dataset as two independent batches for the purpose of batch correction. Cells expressing fewer than 200 genes and genes found in less than 3 cells were filtered out. We normalized the total counts per cell to 10,000 and applied a log1p transformation to alleviate the effects of data sparsity. To improve the signal-to-noise ratio in these datasets, we selected 2000 highly variable genes for further computational analysis. After completing data preprocessing, we performed multimodal batch correction in the same manner as the integration of multiple batches of scRNA-seq data.

### Batch benchmark generation

In order to generate scRNA-seq data with real batch information, scCobra first trains a model by integrating scRNA-seq data from multiple batches. The trained encoder is capable of removing batch effects, while the decoder, equipped with a DSBN layer, can restore batch information, each BN (Batch Normalization) layer within the DSBN layer is used to restore the batch information for a specific batch. At this point, by selecting the source of the batch information for the generated data, the specific BN layer for that batch in Decoder can be activated. Consequently, the decoder can generate single-cell data with the specific batch information.

### Benchmarking with other methods

In our benchmarking analysis, we focused on evaluating a series of computational tools for their effectiveness in the integration and analysis of single-cell transcriptomic data. Our evaluation highlighted their capabilities in addressing batch effects, facilitating data integration, and improving interpretability. (1) Seurat (v3.2.3): Employed for data normalization and the selection of 2000 highly variable genes using the ‘vst’ method, allowing for precise identification of significant features across batches. (2) scVI (scvi-tools, v0.11): Utilized for its advanced batch correction techniques within a Variational Autoencoder (VAE) model by adjusting the ‘n_batch’ parameter, demonstrating its prowess in minimizing batch discrepancies. (3) Harmony (Harmony-pytorch v1.0) and Scanpy: These tools were combined for cell and gene filtering, with Harmony applied after PCA to harmonize datasets, ensuring seamless integration. (4) Scanorama (v1.6): Integrated with Scanpy for merging datasets, it employed the ‘Seurat’ method for selecting highly variable genes, creating an integrated analysis matrix essential for downstream analysis. (5) SCALEX (v1.0): Chosen for its ability to integrate multiple batches of datasets through label transfer functions post-training, SCALEX is crucial for maintaining biological coherence. (6) TOSICA (v1.0.0): Utilized for its direct annotation capabilities using a reference atlas, enabling accurate identification of cell types in unlabeled data batches, thus enhancing the interpretability of the results. Through this exercise, we aimed to provide a comprehensive evaluation of each tool’s performance, shedding light on their utility and effectiveness in single-cell data integration and harmonization.

In this study, we leverage the following metrics to evaluate the performance of all benchmarking methods. For batch correction evaluation, we use the Adjusted Rand Index (ARI), Normalized Mutual Information (NMI), and average silhouette width across cell types (CellType_ASW) to measure the biological conservation of the data. The batch correction performance is evaluated by the following metrics: average silhouette width (Batch ASW) across batches and Batch Entropy Mixing Score. The Adjusted Rand Index (ARI)^69^ serves as a metric for quantifying the similarity between two clustering results, mitigating the probability factors inherent in the Rand Index (RI) during random assignment. The Normalized Mutual Information (NMI)^69^ auges the similarity between two clusters based on mutual information, quantifying the shared information between them. CellType_ASW^70^ measures the relationship between the compactness within a cell clustering and the separation between cell clusters. For batch correction metrics, Batch_ASW^70^ calculates the silhouette coefficient for different batches to assess whether cells from various batches are uniformly mixed within the dataset. Batch Entropy Mixing Score^25^ is a quantitative metric designed to evaluate the homogeneity of cell distribution from different batches in single-cell transcriptomic data. Detailed information on these metrics can be found in the supplementary materials.

## Data availability

In this study, we used one simulated and two actual single-cell datasets to evaluate our method, similar to the datasets that were used in the scIB’s method^53^. Among them, the simulated dataset was generated by the Splatter package^65^. The dataset was composed of 12097 cells with 9979 genes, a mixture of seven cell types from six batches. The pancreas dataset was composed of 14 cell types mixed across nine sequencing platforms, where each sample of InDrop-seq data was regarded as an independent batch, including 16382 cells with 19093 genes, this dataset was collected from Gene Expression Omnibus (GEO) (GSE81076^46^, GSE85241^45^, GSE86469^45^, GSE84133^47^, GSE81608^47^). The third immune atlas dataset contains 10 batches with 16 cell types, including 33506 cells with 15148 genes. This dataset is available with the GEO accession number GSE115189^36^, GSE128066^37^, GSE94820^38^ and https://support.10xgenomics.com/single-cell-gene-expression/datasets/3.0.0/pbmc_10k_v3. Multi-omic data are from https://support.10xgenomics.com/single-cell-gene-expression/datasets/3.0.0/pbmc_10k_v3 and https://support.10xgenomics.com/single-cell-atac/datasets/1.0.1/atac_v1_pbmc_10k. The MERFISH dataset has collected data on 64,373 cells with 155 genes, and the scRNA-seq dataset includes 30,370 cells with 21,030 genes^41^. The datasets use for label transfer evaluation come from GSE83139^71^, GSE114297^72^ and GSE81547^73^. Covid-19 data used in over-correction evaluation is from https://covid19cellatlas.org/^39^

## Software availability

The implementation of scCobra is available at https://github.com/mcgilldinglab/scCobra.

## Acknowledgements

This work was partially supported by CIHR PJT-180505, FRQS 295298, 295299, and NSERC RGPIN-2022-04399 to J.D. Y.X is supported by grants from the National Science Foundation of China (62172274).

