## Supplementary for "scCobra: Contrastive cell embedding learning with domain-adaptation for single-cell data integration and harmonization"

### Supplementary Method

#### Batch Correction and Biological Conservation metrics

**Adjusted Rand Index[1]:** Adjusted Rand Index (ARI) reflects the degree of overlap between the two divisions. This metric requires the true category labels for the data. We use the Leiden clustering method to obtain the predicted labels of the cells. The ARI measures the consistency between the true category labels and the model-predicted labels. The value of ARI is within [0,1], and a larger ARI indicates a higher consistency between the predictions and the ground truth.

**Normalized Mutual Information[2]**: Normalized Mutual Information (NMI) standardized mutual information is commonly used in clustering to measure the similarity of two clustering results. It can objectively evaluate the accuracy of a community division compared with a standard division. The value range of NMI is 0 to 1, and the higher the value, the more accurate the division.

**Average Silhouette Width[3]**: Average Silhouette Width (ASW) across cell types. The silhouette coefficient is a metric to evaluate the quality of clustering. For each cell in the cluster, the best value is 1 and the worst value is -1. A value close to 0 indicates overlapping clusters. A negative value usually indicates that the sample has been assigned to the wrong cluster because different clusters are more similar. Cell-type ASW is a modified approach to measure batch mixing. It computes the silhouette score on cells with identical cell labels and is scaled to a value between 0 and 1. ASW across batches. Batch ASW refers to the Modified ASW of a batch which measures batch correction by calculating the silhouette score of cells in a given batch. Due to the averaging process, the value range is 0 to 1.

**Batch_mix_score**[4]: The Batch Entropy Mixing Score is a metric employed to assess how Batch Entropy Mixing Score is a quantitative metric designed to evaluate the homogeneity of cell distribution from different batches in single-cell transcriptomic data. The computation of this score is methodically structured into several steps. It begins with the determination of cell proportions in each batch relative to the overall cell population. This is followed by the random selection of a subset of cells across batches and the identification of their nearest neighbors to assess local batch mixing. To account for batch size discrepancies, a correction term is applied. The regional mixing entropy is then calculated, encapsulating the dispersion of cells from different batches within the data manifold. By repeating this process across multiple iterations with varying cell selections and averaging the entropy values, the final Batch Entropy Mixing Score is obtained, providing a robust measure of batch integration efficacy within the dataset.

### Supplementary Tables

**Table S1**. Biological conservation and batch correction metrics of five methods on simulation dataset.

| Methods | Biological conservation metrics | | | Batch correction metrics | |
|---|---|---|---|---|---|
| | Cell type ASW | NMI | ARI | Batch ASW | Batch_mix_score |
| Seurat[5] | 0.9 | 0.98 | 0.98 | 0.9 | 0.5 |
| Scanorama[6] | 0.87 | 1 | 1 | 0.91 | 0.27 |
| scVI[7] | 0.85 | 1 | 1 | 0.92 | 0.6 |
| Harmony[8] | 0.88 | 0.97 | 0.97 | 0.89 | 0.41 |
| scCobra | 0.93 | 0.99 | 0.99 | 0.96 | 0.62 |

**Table S2**. Biological conservation and batch correction metrics of five methods on pancreas dataset.

| Methods | Biological conservation metrics | | | Batch correction metrics | |
|---|---|---|---|---|---|
| | Cell type ASW | NMI | ARI | Batch ASW | Batch_mix_score |
| Seurat | 0.68 | 0.91 | 0.9 | 0.85 | 0.54 |
| Scanorama | 0.63 | 0.85 | 0.77 | 0.9 | 0.34 |
| scVI | 0.67 | 0.89 | 0.88 | 0.86 | 0.47 |
| Harmony | 0.67 | 0.9 | 0.9 | 0.89 | 0.53 |
| scCobra | 0.7 | 0.88 | 0.87 | 0.91 | 0.5 |

**Table S3**. Biological conservation and batch correction metrics of five methods on Immune dataset.

| Methods | Biological conservation metrics | | | Batch correction metrics | |
|---|---|---|---|---|---|
| | Cell type ASW | NMI | ARI | Batch ASW | Batch_mix_score |
| Seurat | 0.71 | 0.87 | 0.83 | 0.87 | 0.52 |
| Scanorama | 0.69 | 0.86 | 0.8 | 0.90 | 0.32 |
| scVI | 0.72 | 0.89 | 0.87 | 0.89 | 0.52 |
| Harmony | 0.72 | 0.88 | 0.87 | 0.89 | 0.46 |
| scCobra | 0.75 | 0.89 | 0.88 | 0.92 | 0.52 |

**Table S4**. Biological conservation and batch correction metrics of five methods on three datasets (average).

| Methods | Biological conservation metrics | | | Batch correction metrics | |
|---|---|---|---|---|---|
| | Cell type ASW | NMI | ARI | Batch ASW | Batch_mix_score |
| Seurat | 0.55 | 0.72 | 0.61 | 0.87 | 0.53 |
| Scanorama | 0.57 | 0.74 | 0.62 | 0.91 | 0.34 |
| scVI | 0.63 | 0.79 | 0.72 | 0.88 | 0.5 |
| Harmony | 0.6 | 0.78 | 0.74 | 0.9 | 0.45 |
| scCobra | 0.63 | 0.81 | 0.78 | 0.9 | 0.45 |

**Table S5**. Integration results of five methods on multi-omic dataset.

| Method | Batch Entropy | Silhouette | ARI | NMI | cell type ASW |
|---|---|---|---|---|---|
| Seurat | 0.41 | 0.28 | 0.77 | 0.8 | 0.76 |
| Harmony | 0 | 0.06 | 0.47 | 0.71 | 0.76 |
| Scanorama | 0.01 | 0.07 | 0.55 | 0.71 | 0.8 |
| scVI | 0.47 | 0.26 | 0.76 | 0.75 | 0.76 |
| scCobra | 0.5 | 0.24 | 0.82 | 0.79 | 0.77 |

**Table S6**. Integration results of five methods on spatial / scRNA-seq dataset.

| Method | Batch Entropy | Silhouette | ARI | NMI | cell type ASW |
|---|---|---|---|---|---|
| Seurat | 0.38 | 0.26 | 0.73 | 0.59 | 0.63 |
| Harmony | 0.42 | 0.22 | 0.47 | 0.66 | 0.61 |
| Scanorama | 0.18 | 0.18 | 0.67 | 0.49 | 0.59 |
| scVI | 0.43 | 0.2 | 0.61 | 0.71 | 0.6 |
| scCobra | 0.52 | 0.32 | 0.76 | 0.73 | 0.66 |

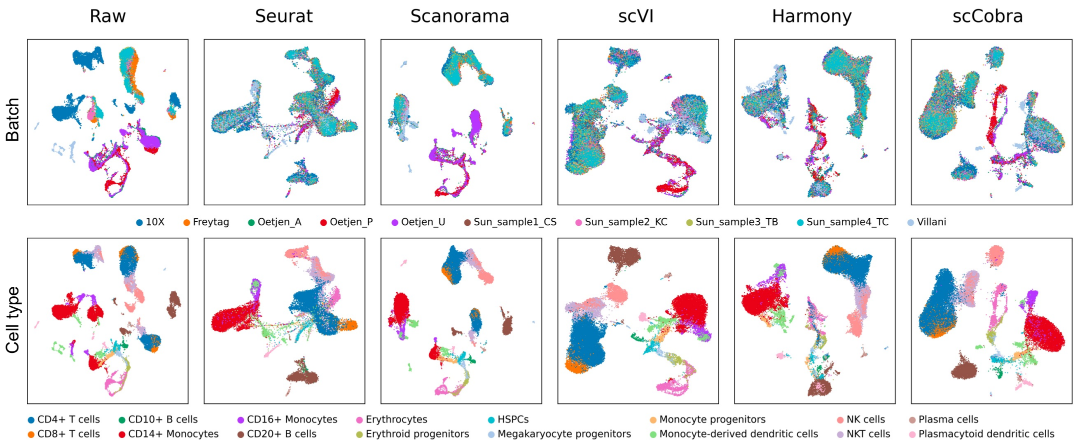


**Fig. S1** UMAP visualization comparing scCobra's performance against benchmarking methods (Seurat, Scanorama, scVI, and Harmony) on the Immune dataset. The visualization is split into two rows: the top row shows batch correction results, with different batches distinguished by color, and the bottom row presents cell type aggregation, with each cell type assigned a unique color.

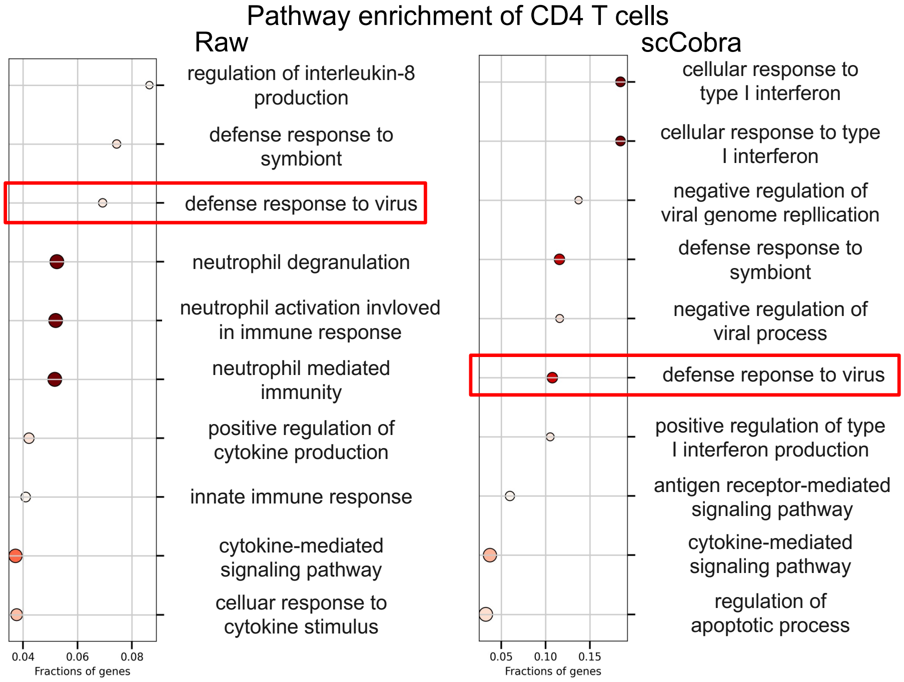


**Fig. S2** Post-correction Gene Ontology enrichment analysis for the COVID-19 dataset underscores scCobra's capability to retain the "Defense response to virus" GO term, which is a top-identified term in the raw (uncorrected) COVID data, crucial for COVID sample analysis. This stands in contrast to other methods, which exhibit correction inaccuracies, evidenced by the loss of this critical GO term, indicative of over-correction and information loss.
